# Evaluation of the Therapeutic Potential of Oral Phycocyanin-Rich Spirulina Extract in Neuropsychiatric Disorders

**DOI:** 10.1101/2021.10.11.463869

**Authors:** Anna Donen, Tzuri Lifschytz, Gilly Wolf, Hagar Ben-Ari, Amit Lotan, Leonard Lerer, Bernard Lerer

## Abstract

**Aim:** Spirulina is a microalga that is widely used as a food supplement and is regarded as having performance enhancing and health promoting properties. We conducted a preliminary evaluation of the possible antidepressant, anti-anxiety, pro-socialization and cognition-enhancing effects of Spirulina in mouse models.

**Methods:** Sixty male BalbC mice aged 3 weeks were administered phycocyanin-rich Spirulina extract (PRSE, 545 mg/kg), fluoxetine (20 mg/kg) or water orally for 5 weeks. During the last 2 weeks of the experiment a series of behavioral-cognitive tests was performed to evaluate motor activity, antidepressant and anti-anxiety effects, socialization and cognitive effects. Effects of PRSE and fluoxetine were compared to those of water.

**Results:** There was a significant effect of PRSE in the activity domain, manifesting as an increase in velocity in the open field (p=0.0007 vs. water). Fluoxetine significantly enhanced immobility in the tail suspension test and the forced swim test reflecting the known antidepressant effect of this compound, but not PRSE. There were no significant effects of PRSE in tests of anxiety, socialization or cognition.

**Conclusions:** The most striking observation in this study was that PRSE significantly enhanced activity in the open field test. Further studies are indicated to confirm and extend this finding and investigate possible mechanisms of action. The results of the current study do not support sporadic reports of possible antidepressant or cognition-enhancing effects of PRSE. Nevertheless, additional studies are indicated using depression models rather than naïve mice, alternative mouse strains, using additional cognitive tests, and administering higher PRSE doses.

## 1. INTRODUCTION

Spirulina is the generally used term for the dried biomass of the organism, *Arthrospira platensis*, a filamentous cyanobacterium with typical prokaryote cell organization (1). Found in alkaline brackish or saline waters in tropical and subtropical regions, Spirulina is most widely distributed in Africa and Asia. Use of Spirulina as food was reported as far back as the sixteenth century in what is now Mexico and in Africa, mainly in the Lake Chad region (2). Spirulina is listed by the US Food and Drug Administration (FDA) under the Generally Recognized as Safe (GRAS) category and is widely consumed as a functional food, especially in Asia, due to its rich content of minerals, proteins, and vitamins (3)

Spirulina has been reported to be an effective health supplement in several studies. A study in Cameroon demonstrated a significant reduction in total cholesterol, triglycerides and low density lipoproteins among HIV positive patients taking Spirulina (versus control) (4). Spirulina has also been reported to reduce body mass and BMI in obese individuals (5) as well as to have possible anti-viral properties demonstrated by its reported ability to inhibit influenza A virus plaque formation and replication. (6) In addition, Spirulina may be able to act as an anticoagulant and reduce platelet activity by reducing platelet volume (7). The anti-inflammatory and antiviral properties of Spirulina extracts are well documented, but their mechanism of action remains unclear. The main active constituent of the Spirulina extracts is phycocyanin, an assembly of phycobiliprotein monomers composed of α and β subunits and their respective chromophores linked via thioether bonds. Phycocyanin has three phycocyanobilin (PCB) chromophores attached to each αβ monomer, which combine to form the typical (αβ)6 hexameric structure of the phycobiliprotein (8). Phycocyanin-rich extracts of Spirulina and Afazononinom flos-aquae (AFA or Klamath Lake algae) are currently available and are marketed as health supplements or “superfoods” with a wide range of purported health benefits (https://www.unicornsuperfoods.com/products/100-natural-blue-spirulina-powder)

The median lethal dose of Spirulina is >5000mg/kg; up to 800mg/kg of body weight have been given to rats orally and up to 2000mg/kg body weight per day subcutaneously with no adverse reaction. No hepatotoxic, cardiotoxic or nephrotoxic or detrimental effects on fertility have been found, nor has Spirulina been noted to be teratogenic (2). Spirulina is widely regarded as one of the safest health supplements and its potential benefits are mentioned by many leading, reputable providers of consumer health advice (9)

Research on Spirulina’s neuropsychiatric and behavioral effects is more preliminary, particularly in humans. Johnson et al reported that exercise output in a small group of untrained men, half of whom received Spirulina (3g/day, orally), measured as number of kcal expended in 30 minutes in a cross-training exercise, increased significantly after one week when compared with control. Spirulina was also associated with decreased mental fatigue in simple mathematical calculations throughout the eight week period of supplementation as well as with an improved subjective state of mental health (10).

In animal models, Spirulina was reported to reverse MSG-induced memory deterioration in rats (11). In a group of mice displaying a phenotype of accelerated aging (12), administration of Spirulina significantly reduced amyloid-beta accumulation in the hippocampus and whole brain as well as decreasing lipid peroxides in the hippocampus and striatum (13). In another study, rats administered Spirulina and exposed to the neurotoxin, 6-hydroxydopamine, showed a relatively diminished reduction in dopaminergic neurons compared to controls (14). A reduction in fluoride-induced anxiety was seen in offspring of rats exposed to Spirulina during pregnancy as shown by increased time in open arms in the elevated plus maze and increased mobility in the open field test (15). Mice fed hydrolysed Spirulina were reported to show significantly decreased immobility in the forced swim test (16). Spirulina may also protect against ischemic damage - when mice were fed Spirulina pellets prior to stroke-inducing surgery, total distance traversed on post-operative behavioral tests was greater. Post-mortem dissection showed decreased infarction size compared to the other groups (17).

To the best of our knowledge, no systematic work has been undertaken to investigate the possible activity-enhancing, cognitive, antidepressant, anti-anxiety and pro-socialization effects of Spirulina. The present study was undertaken to systematically investigate the possible effects of Spirulina in these domains as a basis for clinical studies. We performed a preliminary study using a phycocyanin-rich Spirulina extract (PRSE) prepared by Algonesia Technologies SAS, France. PRSE was chosen for study because phycocyanin-rich Spirulina extracts (blue Spirulina) are growing in popularity among nutritional supplement consumers. This trend is mainly driven by the convenience of being able to add smaller amounts to smoothies and foods as compared to Spirulina powder (less than half a teaspoon versus 2-4 tablespoons), which has a bitter taste and unappealing colour. We used a series of behavioural and cognitive tests to study the effect of PRSE administered to mice in drinking water for three weeks.

## 2. Methods

### 2.1 Animals

Sixty male BalbC mice aged 8 weeks were used in this study (supplied by Envigo Labs, Israel). Mice were housed in the SPF-certified facility of the Sharett Institute, Hadassah-Hebrew University Medical Centre. Food and water were provided *ad libitum*. Mice were kept under a 12-hour dark/light cycle (lights on at 7am). Mice were weighed twice weekly from the beginning of treatment. The experiments were approved by the Hebrew University Ethics Committee on Animal Care and Use (Application MD-17152604). As an American Association for Accreditation of Laboratory Animal Care (AAALAC) accredited institute the Hebrew University Ethics Committee follows the National Research Council’s (NRC) “Guide for the Care and Use of Laboratory Animals”.

### 2.2 Procedures

Mice were divided into 3 treatment groups, 20 mice in each group, each receiving different drinking solutions. Mice entered the experiment in a staggered fashion in three cohorts, approximately 20 animals in each cohort with 6-7 animals in each of the three different treatment groups. The control group received regular drinking water; the positive control group received fluoxetine at a concentration of 20mg/kg body weight and the treatment group received phycocyanin-rich Spirulina extract (PRSE; Algonesia Technologies SAS) at a concentration of 545mg/kg body weight (phycocyanin concentration 30%, as measured by the method of Bennett and Borograd (15) Following a pilot study, mice were assumed to drink 3.5ml water per day. Three weeks after the beginning of the experiment, while still receiving the treatments, animals underwent a behavioural-cognitive test battery. A timeline is shown in Figure 1.

**Figure 1:**
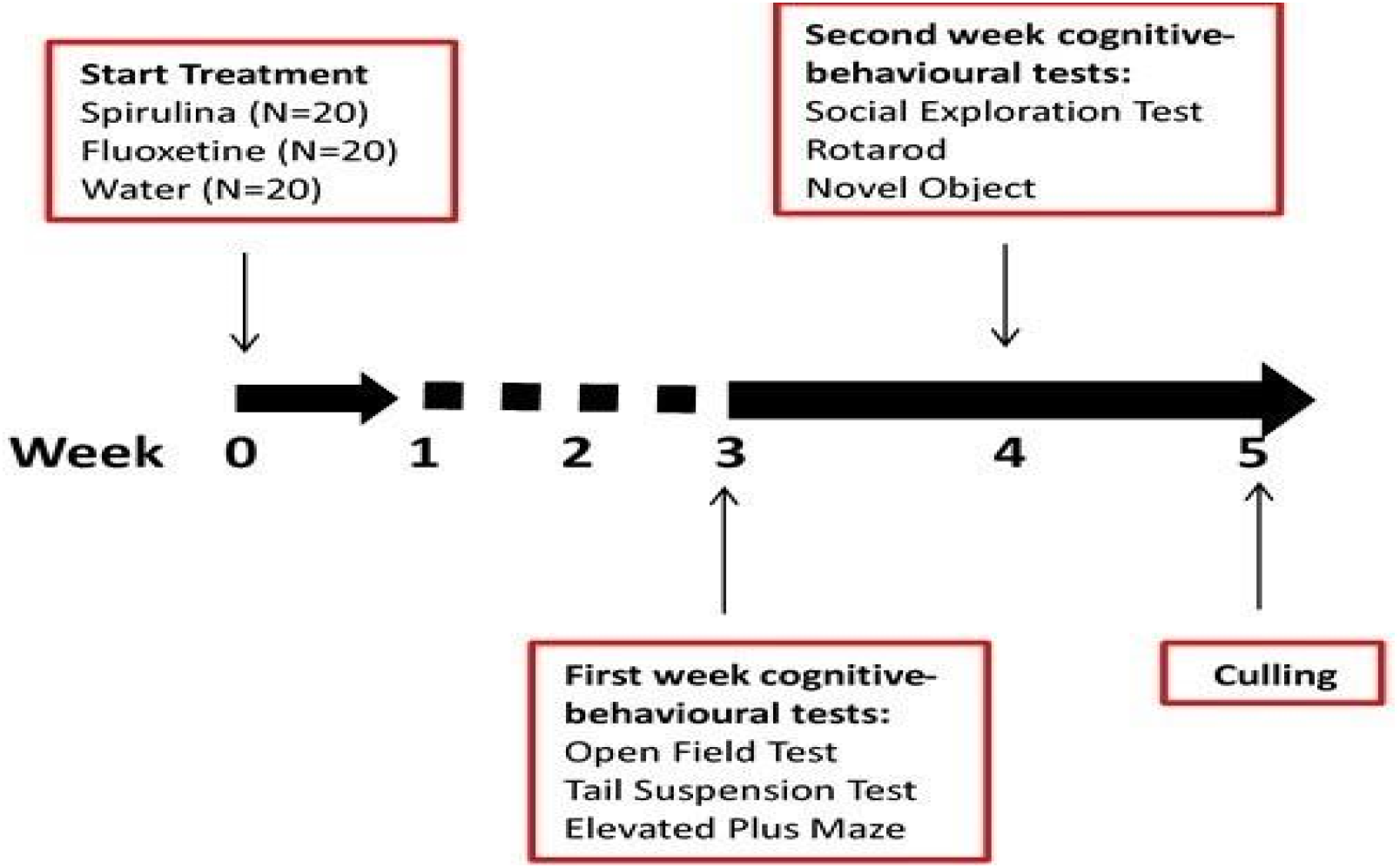
Timeline of procedures; (N=number)

### 2.3 Behavioral-Cognitive Tests

Mice underwent a 2-week behavioral-cognitive test battery; most were recorded and analysed using the Ethovision 11 system (Noldus Information Technology Inc.). When scoring was done manually, the scorer was blind to the treatments received by the mice. Tests were performed in the same order for each of the three treatment groups.

#### Open field test (OFT)

Mice were placed in a 50 × 50 cm arena surrounded by 40 cm walls for a 6 min test. The centre of the arena was defined as a 25 × 25 cm square in the middle of the arena. Velocity of movement is the primary outcome and is a test of motor skill. Secondary outcomes are number of entries made to the centre of the arena and time spent in the centre of the arena. The open field test was performed during the first week of the behavioral battery, using the Ethovision 11 system, providing fully computerized, blinded and unbiased measurement.

#### Tail suspension test

For 6 minutes mice are suspended upside down by adhesive tape placed 1 cm from the tail tip, at an elevation of 50 cm from the nearest surface. Every 5 seconds the animal’s behavior is noted to be “mobile” or “immobile” by a blinded observer. Magnitude of immobility is compared between different treatment groups.

#### Forced swim Test

Animals are individually placed in a 21 (diameter)x 46 (height) cm circular transparent plexiglass tank filled with 15 cm of room-temperature water for 6 min. Over the final 4 min, the time in which mice are actively swimming (mobility) and the time of passive floating (immobility) is recorded by the Ethovision system and scored by a blinded observer. Anti-depressant medication has been shown to decrease immobility in the tail suspension and forced swim test (16,17)

#### Elevated plus maze

The test apparatus consists of two open arms (30&5 cm) bordered by a 1 cm high rim across from each other and perpendicular to two closed arms bordered by a rim of 16 cm. The centre of the maze, in which the 4 arms converge, is a 5&5 cm platform. The entire maze is lifted 75 cm from the floor. Mice were placed in the central platform of the maze and were allowed to explore it for 5 minutes. Durations and number of visits in both the open and closed arms were recorded as a measure of anxiety. This test was recorded using the Ethovision system.

The rotarod, social interaction and novel object recognition tests were also carried out and are described in the Supplementary Information.

### 2.4 Statistical analysis

Data were analysed using SPSS 24. One-way analysis of variance (ANOVA) was performed, with repeated measures when appropriate, followed by planned comparisons. Analysis of covariance (ANCOVA) was performed when a significant baseline difference was demonstrated. P values <0.05, two tailed, were regarded as statistically significant.

## 3. Results

### 3.1 Weight

All animals gained weight during the experimental period with no significant differences in weight gain from baseline across the groups.

### 3.2 Volume of liquid drunk

Mice in the fluoxetine group drank significantly less than those in the water group (2.45±0.64 ml. versus 3.70±0.34 ml) (ANOVA: p=0.01, planned comparisons: p =0.01). There was no difference between mice treated with PRSE and those that received water only in the volume of liquid drunk.

### 3.3 Behavioral-cognitive tests

#### 3.3.1 Activity

##### Open field test (OFT)

Administration of PRSE caused a significant increase in velocity of the mice in the open field (one-way ANOVA (*F*(2,59) = 8.122, *p*= 0.001; planned comparisons: PRSE versus water, p = 0.007 fluoxetine vs. water, p= 0.274. Figure 2ii) shows these results graphically and figure 2i) shows a heat map which represents the amount of movement of representative mice from the water (left) and PRSE (right) groups in the arena. Mice receiving PRSE entered the centre of the open field arena significantly more frequently than other animals (one-way ANOVA (*F*(2,59) = 3.869, *p* = 0.027). However, this effect was related to velocity and when the results were analysed with velocity as a covariate, the effect was found to be non-significant. No significant effect of PRSE or fluoxetine was found on duration of time spent in the centre of the arena.

**Figure 2:**
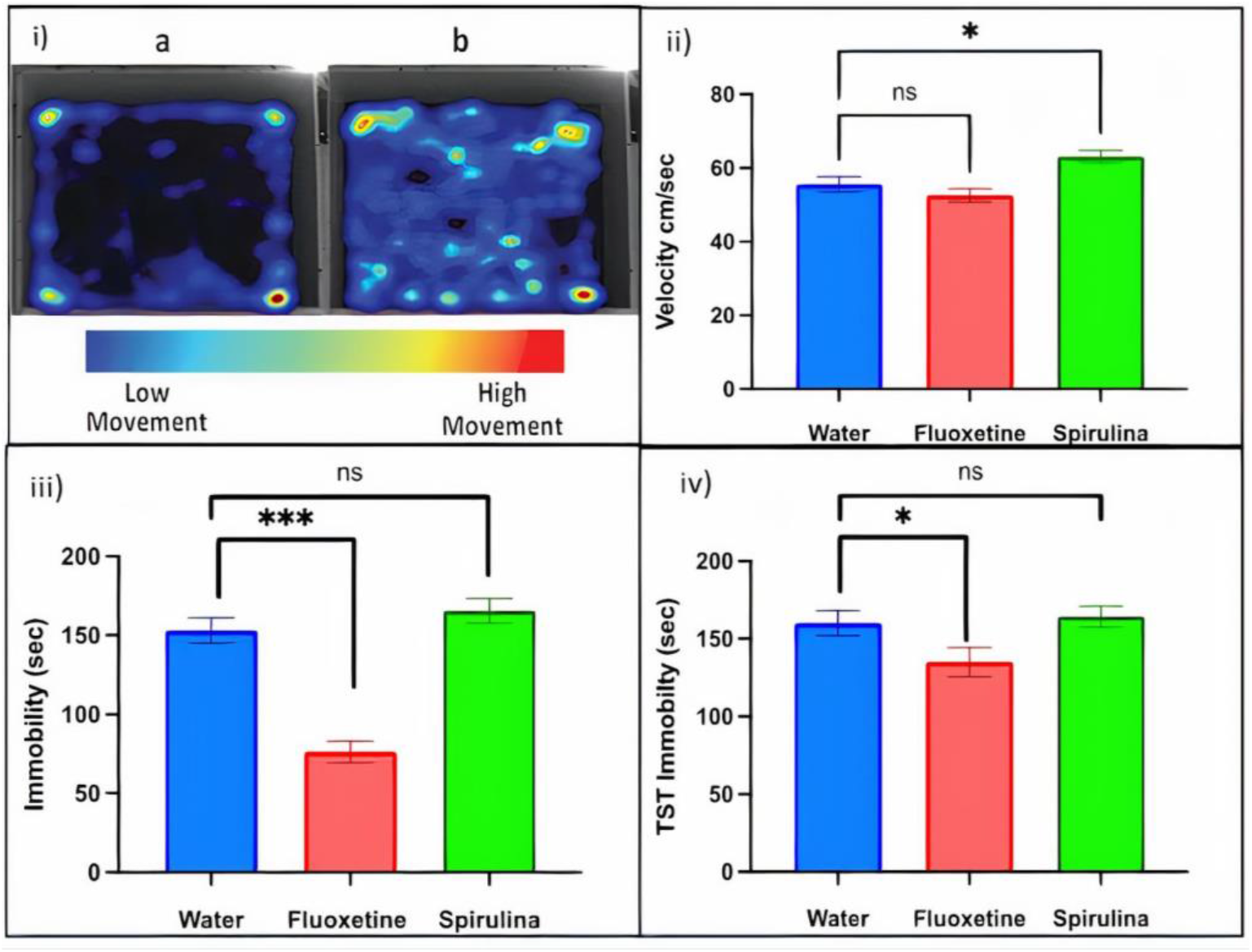
i) Heat maps of mouse movement in the open field test showing (a) increased movement of representative mice receiving water (b) compared to PRSE. ii) velocity during the OFT. Bars represent mean+SEM. &p<0.05. iii) FST results. Bars represent mean +/− SEM time spent immobile. &&p<0.0001. iv) Tail suspension test results. Bars represent mean +/− SEM time spent immobile. &p<0.0

#### 3.3.2 Antidepressant-like effects

##### Forced swim test (FST)

There was a significant reduction in immobility in animals administered fluoxetine versus water and spirulina (one-way ANOVA F (2,57) = 40.89, p<0.0001; planned comparisons: fluoxetine vs water, p<0.0001 but not in the immobility of mice administered PRSE (p=0.4941). See Figure 2iii).

##### Tail suspension test (TST)

There was a significant reduction in immobility in animals administered fluoxetine versus water (one-way ANOVA F(2,59)=3.734, p=0.03; planned comparison: fluoxetine vs. water, p=0.035); but not in the immobility of mice administered PRSE (p= 0.715). See Figure 2iv).

#### 3.3.3 Anxiolytic-like effects

##### Elevated plus maze

No differences were seen between the groups in time spent in the open arms (one-way ANOVA (F(2,59) = 0.091, p = 0.913) or number of entries made to open arms (one-way ANOVA (F(2,59) = 0.145, p = 0.865).

The Rotarod (motor strength), Social Exploration (social interaction) and Novel Object Recognition (perceptual memory) tests did not show significant differences between the experimental groups (Data available on request)

## Discussion

We have reported the effect of chronic, oral administration of phycocyanin-rich Spirulina extract (PRSE) on the performance of BalbC mice on behavioral and cognitive tests relevant to potential therapeutic effects in neuropsychiatric disorders.

The most striking effect observed was in the motor domain – an increase in velocity in the open field test. Animals receiving PRSE moved significantly faster in the arena than those receiving water. Due to their increased velocity, PRSE-treated mice made a significantly greater number of entrances to the centre of the arena compared with other mice. Positive motor findings were not found in the rotarod test which reflects muscle strength and coordination.

While preliminary, the effects of PRSE on activity in the open field that we observed are noteworthy. They raise the interesting possibility that Spirulina could have a role as a performance-enhancing substance in sports. Further studies are indicated to replicate this finding and to determine whether it is evident in other paradigms that measure motor performance in rodents. Our finding that rotarod performance is not enhanced suggests that PRSE does not affect muscle strength and coordination. The significant effect of PRSE on velocity in the open field test is of considerable interest in view of the widespread use of Spirulina as a booster of athletic performance (http://www.algae.company/spirulina-for-athletes/). There is growing interest among consumers, especially those actively engaged in sports and online gaming, in natural products that enhance both physical and cognitive performance. There is also a growing demand for “clean label” formulated sports drinks that have a limited number of effective ingredients, are low in sugar and provide some proteins (https://www.naturalproductsinsider.com/healthy-living/sports-nutrition-beverages-redefined-download). A comparison of PRSE with the effects of non-prescription, performance enhancing substances that are available on the market, such as caffeine and taurine, is indicated. If the finding is further supported in additional experiments, studies aimed at elucidating the underlying biological mechanisms should be performed. Substances such as caffeine and taurine are associated with physiological side effects such as tachycardia and hypertension as well as potential nephrotoxicity (18). Thus, the possibility that Spirulina may have similar or greater benefits but less or no side-effects calls for further investigation.

There was no significant effect of PRSE on immobility in the forced swim test or tail suspension test; this does not support a potential antidepressant effect. This is contrary to a previous observation in which PRSE was found to decrease immobility in the forced swim test. (16) In the current experiment, reduced immobility on the tail suspension test and forced swim test was induced by fluoxetine, a widely used antidepressant that has previously been shown to reduce immobility on this test in BalbC mice on oral administration (19). This finding suggests that PRSE, at the dose studied by us, does not have significant antidepressant effects. Further studies at a higher PRSE dose are indicated in order to exclude a possible antidepressant effect.

No significant effect of PRSE was seen on behavioral tests assessing anxiety, as reflected by the results of the elevated plus maze and the duration of time spent in the centre of the arena in the open field. There was also no significant effect on social behaviour assessed by the social exploration test, nor on perceptual; memory assessed by the novel object recognition test. This contrast with previous studies may be due to the different cognitive domains that were measured in our experiment by the NOR test compared to the Barnes and active and passive avoidance tests. (14, 16). Our findings do not suggest an effect of PRSE at the dose studied on anxiety, social behaviour and cognitive function. However, further studies at higher doses and using other behavioural paradigms are indicated.

In summary, the results of the current study do not support sporadic reports of possible antidepressant or cognition-enhancing effects of PRSE even though the formulation we used was of greater purity than that used in prior experiments. Additional studies are indicated using depression models rather than naïve mice and a wider range of cognitive tests should be performed. Higher PRSE doses should be administered. Testing the effect of PRSE in mouse strains other than BalbC is also indicated.

It should be noted that in this experiment mice receiving fluoxetine and PRSE drank less than expected on the basis of our pilot study (3.07ml/day as opposed to 4.5ml/day). They therefore received less compound than intended (fluoxetine 12.8mg/kg/day versus 20mg/kg/d and PRSE 464mg/kg/day versus 545mg/kg/day). The fluoxetine group in particular drank significantly less than the water group; nevertheless, the amount of daily fluoxetine consumed was greater than the 10mg/kg in rodents that has been reported to have antidepressant effects (16) and we did see a significant effect of fluoxetine in the TST and FST. The net quantity of PRSE drunk was similar to the dose generally proposed as a nutritional supplement (500mg/kg). It is therefore unlikely that sub-optimal dosing was responsible for the lack of efficacy seen in depression, anxiety and cognitive domains.

## Conclusion

While no significant, effects of PRSE were observed on tests reflecting possible antidepressant, anti-anxiety, pro-social and cognition-enhancing effects, these tests should be repeated at higher PRSE doses, using additional test paradigms and in additional mouse strains. The significant effect of PRSE on velocity in the open field test, although preliminary, is of considerable interest and should be studied further in order to investigate whether Spirulina could potentially have a role as a performance-enhancing substance in sport.

## CONFLICT OF INTEREST

Leonard Lerer held an equity stake in Algonesia Technologies SAS during the period that this study was conducted

## ACKNOWLEDGEMENTS

None

## FUNDING

Supported in part by Algonesia Technologies SAS

## HUMAN AND ANIMAL RIGHTS

The experiments were approved by the Hebrew University Ethics Committee on Animal Care and Use (Application MD-17152604). The experimental procedure was in accordance with the ARRIVE guidelines.

## DATA ACCESS STATEMENT

Access to the data may be obtained by contacting the corresponding author, Prof. Bernard Lerer -

